# Macroscopic control of synchronous electrical signaling with chemically-excited gene expression

**DOI:** 10.1101/2022.01.19.476902

**Authors:** M. García-Navarrete, Merisa Avdovic, S. Pérez García, D. Ruiz Sanchis, K. Wabnik

## Abstract

Excitable cells can convert electrical signals into chemical outputs to facilitate the active transport of information across larger distances. This electrical-to-chemical conversion requires a tightly regulated expression of ion channels. Alterations of ion channel expression provide landmarks of numerous pathological diseases, such as cardiac arrhythmia, epilepsy, or cancer. Although the activity of ion channels can be locally regulated by external light or chemical stimulus, it remains challenging to coordinate the expression of ion channels on extended spatial-temporal scales in a non-invasive manner. Here, we have engineered yeast *S. cerevisiae* to read and convert local chemical concentrations into a dynamic electrical field distributed across cell populations. The core mechanism encodes a chemically-excitable dual-feedback gene circuit that precisely tunes the expression domain of potassium channels, globally coordinating cyclic firing of the plasma membrane potential (PMP). We demonstrate that this mechanism leverages an engineered constitutively open bacterial potassium channel KcsA to directly couple chemical stimuli with ion flux through gene expression and it can interface with the host ion channels through the pulsatile production of toxins. Our study provides a robust synthetic transcriptional toolbox underlying the conversion of local chemical environments into spatiotemporally organized electrical impulses for various cellular engineering, synthetic biology, and potential therapeutic applications.

## Introduction

Communication through electrical signals provides an effective mechanism for the rapid delivery of information across noisy cells and tissues. Prominent examples of this phenomenon across kingdoms include excitable neuronal circuits (1), plant defense signaling (2), and metabolic coordination of biofilm growth (3). These seemingly different forms of electrical signaling involve ion channels (4). Outward-rectifying potassium channels release potassium from the intracellular reservoir to the extracellular space, thereby allowing for potassium exchange between neighboring cells (5). A rapid gating of ion channels at the subcellular level maintains the balance in the plasma membrane potential (PMP), which in turn provides power reservoir, and electrical communication routes that are central to cell physiology (6).

The activity of potassium channels can be locally modulated by voltage-gating, mechanical or light stimulus, and external ligands. In the last decade, advances in optogenetics and chemical biology of ion channels have powered numerous medical applications through local regulation of electrical activity in living cells with the ultimate goal of treating major life-threatening diseases (7–10). Interestingly, recent studies indicate the spatial-temporal regulation of ion channel expression is critical for landmarking pathological conditions such as cardiac arrhythmia, epilepsy, or various types of cancer (11–15). However, a major challenge is to achieve robust control of ion channel expression on extended spatial-temporal scales in a non-invasive manner. Such a strategy would build a foundation for advanced applications in treating epilepsy, chronic pain, irregular heartbeats, or potentially various types of cancer. More broadly, it would create new opportunities for controlling synchronous electrical activity on a macroscopic level in living systems. Yet, a key bottleneck in achieving this goal lies in the inherent noisy expression of ion channels at a single-cell level. To our knowledge, no robust solution has been presented to overcome this limitation.

To address this challenge, we build a synthetic gene regulatory system that is capable of effectively buffering cellular noise, coordinating ion channel expression in the cell populations of the model eukaryote *S. cerevisiae*. This synthetic system builds upon gene circuits that respond to chemical stimuli (16) through coupled Mar-family receptors (MarR and IacR)(17). These circuits can synchronize cell responses by using a rate-dependent hysteresis mechanism (16). Here, we modified these circuits to encode intrinsic excitability (Fig. 1A), thereby linking chemical excitability to robust ion channel expression coordinated at the macroscopic scale. By combining computer modeling with live-cell imaging in microfluidic devices we demonstrate that this strategy is plausible, robust, and suitable for macroscopic control of synchronous electrical signaling in living eukaryotic communities.

**Fig. 1.**
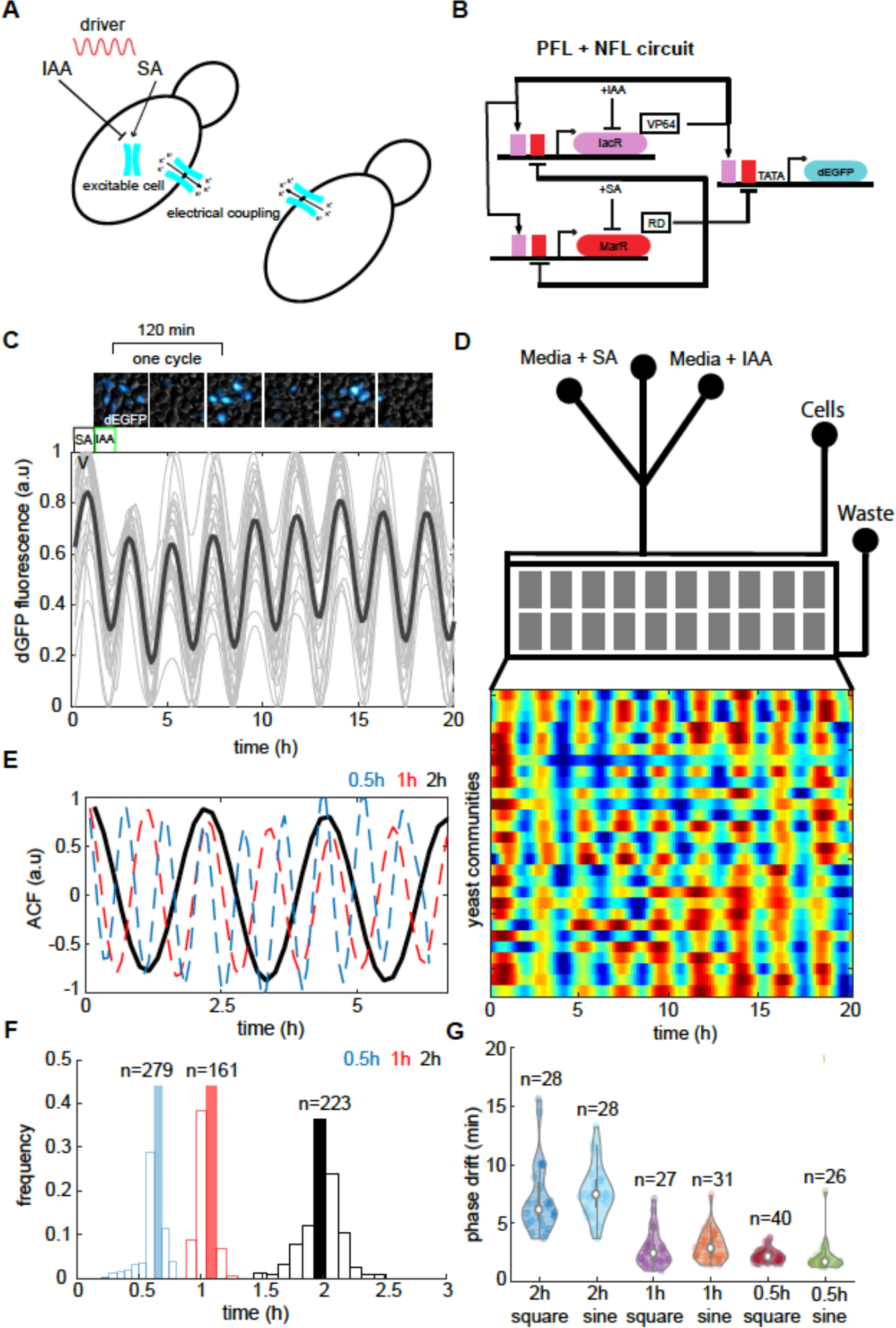
Experimental validation of excitable gene circuit driven by phytohormone clock. **(A, B**) A prototype scheme of the chemically excitable system for tuning ion channel expression. Two phytohormones IAA and SA control the transcription of the potassium channel (cyan) through coupled positive and negative feedback circuits (B). IacR sensor acts as an activator in a positive feedback loop (PFL) and MarR is a repressor in the negative feedback loop (NFL). dEGFP reporter is a short-lived EGFP protein placed under the control of IacR/MarR regulated promoter. Each cell is interfaced with the environment through the detection of IAA-SA stimuli, eventually resulting in the electrical signal coupling within the consortia. **(C)** Top panel, sample screenshots from microfluidic experiments with cells carrying excitable circuits (B). Bottom panel, dEGFP time traces from all traps recorded in the microfluidic device show distinct peaks of dEGFP fluorescence. **(D)** Schematics of the microfluidic device (top panel) with input and output ports and cell trapping regions, and heat map of dEGFP fluorescence for each of recorded trapping regions (n=28 yeast communities) corresponds to time traces(C). Note the full synchrony between individual yeast populations (n=26, 27, 28). (**E**) Cumulative auto-correlation function (ACF) of dEGFP reporter for all recorder population trapping regions recorded for three different periods of IAA-SA environmental stimuli. (**F**) Observed frequencies of periodic dEGFP response. The period of dEGFP oscillations is matched across yeast communities as shown by sharp period distributions. (**G)** Minimal phase drift (∼5%) indicates remarkable coordination across populations of cells, regardless of phytohormone clock period or shape of environmental stimulus.

## Results & Discussion

### Excitable system design

Briefly, our strategy includes a three-component gene circuit that controls potassium channel expression within the *S. cerevisiae* community following dynamic changes of chemical concentrations in the environment (Fig. 1A). We chose Mar receptors (consecutively named MarR and IacR)(16, 17) to facilitate dual-chemical perception of phytohormones salicylate acid (SA) and indole acetic acid/auxin (IAA) in the environment (Fig. 1B). Next, to encode excitability (18) we coupled IacR and MarR through positive and negative feedback loops in the dual-feedback synthetic gene circuit (Fig. 1B). To analyze the robustness and dynamics of the designed circuit we first performed computer model simulations of the IacR-MarR feedback system and identified regimes that demonstrate the prominence for the excitable dynamics (Fig. S1). Model predictions revealed minimal prerequisites for the transit between steady-state (Fig. S1A), excitability (Fig. S1B), and oscillations (Fig. S1C). These conditions directly relate to the ratio of IacR-MarR deactivation (bifurcation point) which depends primarily on phytohormone concentrations in the environment (Fig. S1A-C). Indeed, *in silico* periodic stimulations with IAA and SA predicted the sustained excitable response of the circuit triggered by phytohormones (Fig. S1D).

To test these model predictions, we implemented a synthetic circuit in *S. cerevisiae* by tagging IacR with herpes simplex virus trans-activation domain (VP64) (19) and MarR with repression Mig1 silencing domain (20) (Fig. 1B, Fig. S2). Initially, we tested that this system responds to a broad range of IAA and SA concentrations in high-throughput time-lapse micro-well plate fluorescence assays (Figs. S3 and S4). Next, we employed a controlled environment on the microfluidic chip where cells grow under continuous media perfusion (Fig. 1C and D). We stimulated dozens of engineered yeast populations with antithetic pulses of SA and IAA of either square or sine wave patterns with different frequencies to mimic diverse chemical environments. Throughout the experiment, we could control the period of stimuli with a microfluidic flow control system and record fluorescence changes of unstable dEGFP reporter (21) that reports the outcome of circuit dynamics (Movie S1). Spatially coordinated pulses of dEGFP were observed under phytohormone stimulation as rapidly as 30 minutes (Fig. 1C, D, Fig. S5A-D). The robust unison of gene expression was confirmed further by the analysis of cumulative dEGFP signal autocorrelation (Fig. 1E), frequency response spectrum (Fig. S6A), and period distribution (Fig. 1F). Typically, less than 5% phase shift was observed in the response of individual colonies (Fig. 1G), indicative of global coordination of gene expression among even distal fungi populations in both chemical microenvironments (sine or square-shaped waves) (Fig. S5). Cell populations also exhibit a low dispersion (coefficient of variation (CV) < 1) of dEGFP reporter amplitudes (Fig. S6B), periods (Fig. S6C), and peak widths (Fig. S6D), indicative of general robustness of transcriptional output.

In summary, this chemically-excited circuit presents a promising regulatory module for engineering the coordination of ion channel expression in cell populations.

### Coupling excitable circuit to potassium channel expression at the population level

The plasma membrane potential of yeast is regulated through voltage-gated potassium channels such as the outward-rectifier channel TOK1 (22, 23). TOK1 is the main target for several toxins and volatile anesthetic agents (24), which cause uncontrolled opening and leakage of potassium ions to the extracellular space. Overexpression of TOK1 causes membrane hyperpolarization (more negative PMP) while *tok1* mutation leads to membrane depolarization often accompanied by cell death (25). Thereby, TOK1 controls potassium release from yeast cells and maintains a balance in the PMP.

Next, we sought to wire our excitable circuit (Fig. 1A, B) to the control of PMP firing directly via modulation of potassium channel expression (Fig. 2A). For that, we used the constitutively open KcsA bacterial potassium channel (26, 27) to replace the dEGFP reporter in the output branch (Fig. 2A). This particular strategy bypasses voltage-dependent channel opening allowing the control of PMP solely through the chemically-excited gene expression (Fig. 2A). To confirm the suitability of this strategy, we next measured PMP dynamics in the engineered yeast using a cationic dye Thioflavin T (ThT), previously used to correctly monitor PMP in yeast (28) and bacteria (3). This dye shows a substantial increase in fluorescence following membrane hyperpolarization and its translocation to the cell interior acting as a Nernstian voltage indicator.

**Fig. 2.**
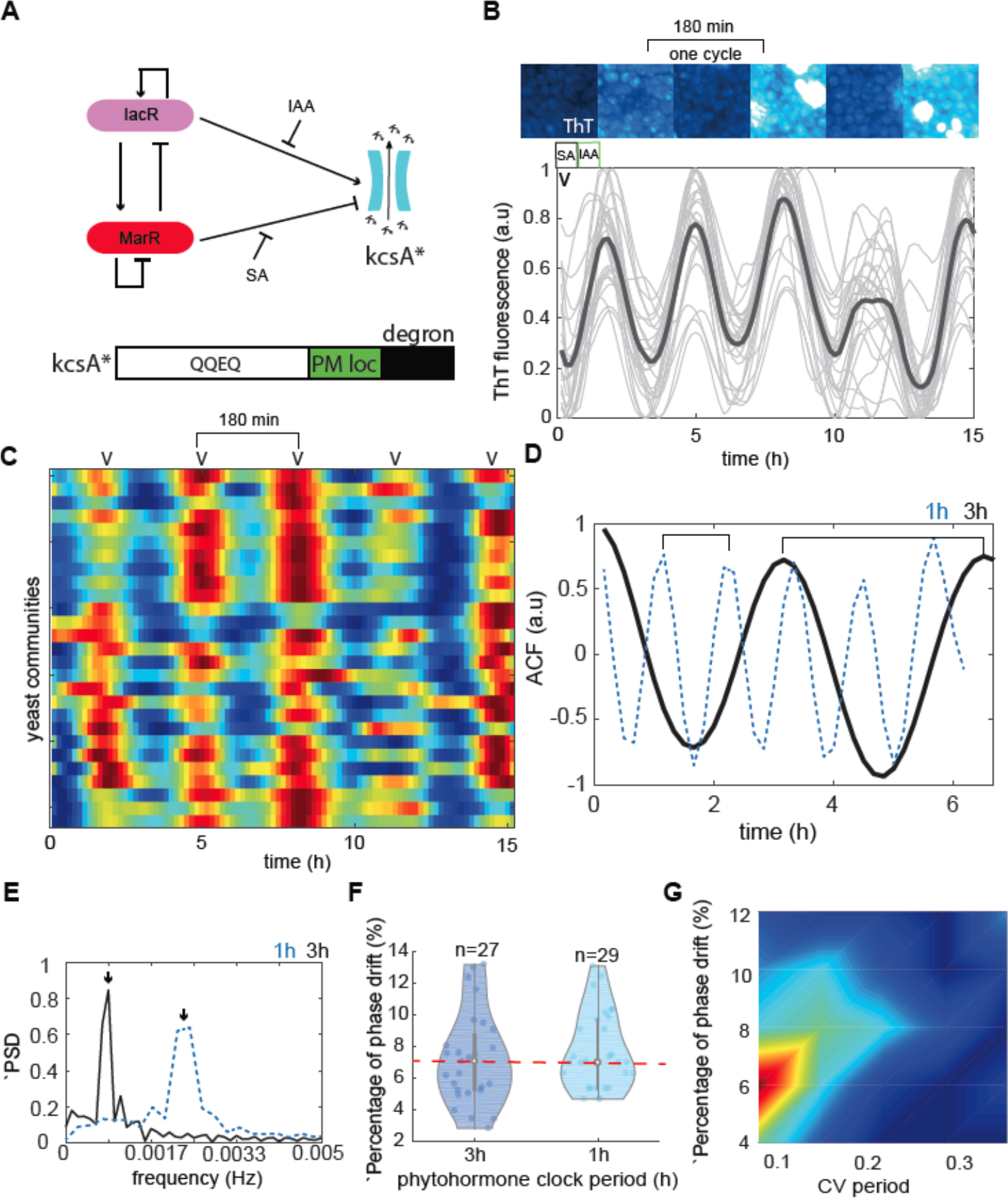
A synthetic dual feedback circuit coordinates globally ion channel expression and electrical firings in yeast communities. **(A)** Schematics of the chemical-electrical converter system contain a dual feedback module (Fig. 1B) that controls the potassium channel activity (KcsA*, lower panel). KcsA* is a constitutively open bacterial potassium channel with plasma membrane localization (PM loc, green) and c-terminus mouse ODC degron (degron, black) sequences. **(B)** Screenshots and time traces from microfluidic experiments display successive cycles of PMP bursts by recording the ThT fluorescence. The period of the IAA-SA environmental clock was set to 3 hours. **(C)** a heat map of ThT fluorescence across all recorded yeast communities corresponds to time traces (B). **(D)** The comparison of cumulative auto-correlation (ACF) of ThT fluorescence for 1 hour (blue dash line) and 3 hours (black line)phytohormone stimulation periods. Note the clear periodic pattern of PMP changes synchronized across populations (**E**) ThT Frequency spectra (PSD) of globally coordinated populations for 1-hour (blue dash line), and 3-hour (black line) phytohormone stimuli. (**F**) The percentage of oscillatory drift as the function of the phytohormone stimuli period. Note that 93% of recorder yeast colonies show the same timing of PMP firings with the speed that is dependent on phytohormone rhythms. (**G**) Density plot of phase drift percentage in the function of ThT period CV. Note a strong correlation between tight coordination of ∼93% yeast communities(red cluster) and low variance of ThT period.

A chemical potassium flux clamp experiment with the graded increase of KCL further confirmed the sensitivity and responsiveness of ThT dye as a PMP marker in yeast cells (Fig. S7).

Remarkably, growing engineered yeast in the microfluidic device subjected to different speeds of chemical stimuli revealed cyclic bursts of PMP (Fig. 2B, C) that were coordinated across cell populations (Fig 2B, C, and Fig. S8A, and Movies S2 and S3). Notably, this type of dynamics was not observed in the control yeast populations subjected to identical environmental fluctuations (Fig. S8B, Movie S4). These data indicate that the chemically-excited ion channel expression is solely responsible for coordinated PMP bursts in engineered yeast populations. The subsequent analysis of period distributions among yeast colonies confirmed a dominant frequency of PMP firing that matched closely that of phytohormone stimuli (Fig. 2C). Similarly, cumulative autocorrelation (Fig. 2D) and frequency spectra (Fig. 2E) analyses confirmed robust coupling of PMP within and across distant populations. Furthermore, the relative variation of amplitudes, as well as the period of PMP firings across colonies, was remarkably low (Fig. S8C-F), indicative of the effective communication of electrical messages within the populations by controlling potassium channel expression at the single-cell level. We also evaluated the percentage of PMP phase drift between individual yeast populations and again we found remarkably very low phase differences (Fig. 2F) that strongly correlate with low variation of the THT period (Fig. 2G and Fig. S8E) confirming further the robust electrical coupling between distant communities.

To further illustrate how coordinated electrical signaling may emerge from chemically- regulated ion channel expression in yeast consortia we developed a spatial-temporal computer model (Fig. S9) that combines the excitable circuit module (Fig. 2A) with Hodgkin–Huxley model of electrical signals(action-potentials) (29). Model simulations of the populations excited by a source cell show the robust propagation of PMP bursts across populations (Fig. S9A). Moreover, the frequency of phytohormone changes in the environment similarly determines the population reaction speed to chemical stimulus by fine tuning potassium channel expression, as observed in our experiments (Fig. S9B). Therefore, experiments and model simulations jointly show how a chemically-excitable cell can propagate the electrical message to its neighbors by dynamically modulating the expression pattern of potassium channels.

### An integration of excitable circuit with the host

We demonstrate that excitable circuit tunes heterologous ion channel expression to generate robust electrical signals coordinated on the macroscopic scale. Next, we asked whether this system can be used to modulate the activity of native potassium channels in the sister population of cells to allow for inter-strain electrical communication. We sought to use a specific K1 toxin that controls the open state of the TOK1 potassium channel (30). The strategy involves regulation of toxin expression level under the control of a synthetic circuit (Fig. 1A) to directly modulate the opening state of the TOK1 channel. For that purpose, we wired our circuit to the transcription activation of the potent K1 killer toxin (Fig. 3A) in the engineered yeast “killer” strain. To test this scenario, we cocultured ‘killer’ (shown in red) with a toxin-sensitive strain in the microfluidic device (Fig. 3B). Release of toxin is controlled via synthetic circuit in response to 2-hour lasting phytohormone stimuli. We monitored ThT fluorescence in the sensitive strain and recorded robust pulses of ThT in the sensitive strain that were coordinated in the sensitive yeast population (Fig. 3C and 3D, Movie S5). A detailed analysis of autocorrelation (Fig. 3E) and phase differences (Fig. 3F) showed that sensitive strain responds to toxin pulses by adjusting PMP firings of each individual cell in the population.

**Fig. 3.**
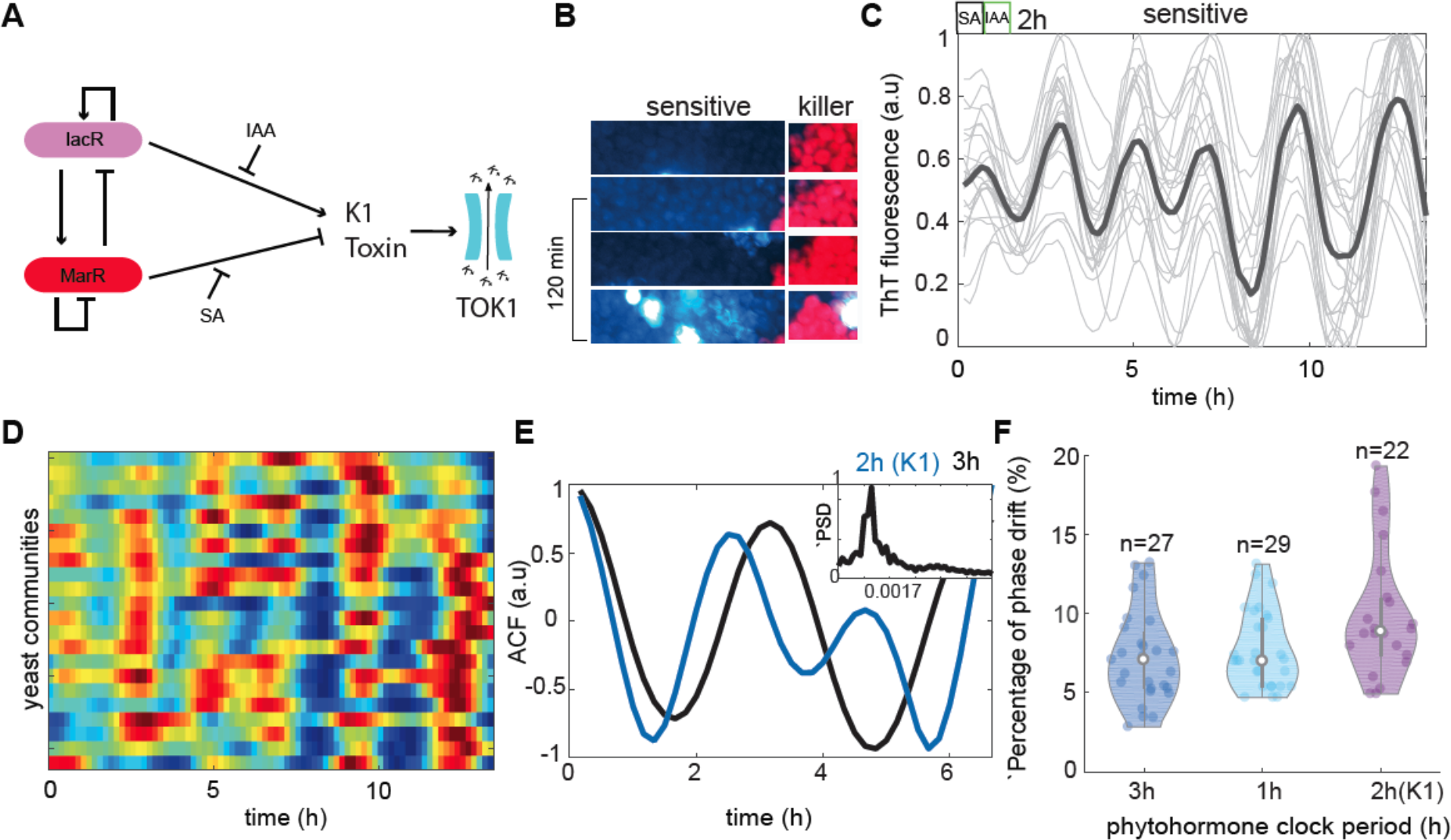
Inter-strain electrical signaling modulated through regulated expression of potassium channel-targeting toxin. **(A)** Schematics of electrical signal modulation via integration with the TOK1 potassium channel. The excitable circuit (Fig. 1B) controls K1 toxin expression. The toxin is released to extracellular space and activates TOK1 channel-dependent potassium flux. (**B**) Example screenshots from co- culture experiments of toxin-producing (mCherry) and sensitive (ThT) strains in the microfluidic device. (**C**) ThT time traces from all traps recorded in the microfluidic device show distinct peaks of PMP potential. (**D**) Heatmap of ThT dynamics in all recorder traps. (**E**) Comparison of cumulative autocorrelation between KcsA* circuits (Fig. 2A) and K1 circuits (A). In both scenarios, a clear oscillatory trend can be observed. The frequency response of the K1-based circuit is presented in the top panel. (**F**) A coordination of PMP firings at inter-community level (phase drift percentage) in the KcsA system (3h, 1h stimuli), and K1-based system (2h stimulus) varies between 90% to 94%.

Thus, our findings demonstrate that electrical activity in the two-strain consortia closely follows the pattern of toxin expression that is controlled by chemical rhythms in the environment.

### Concluding remarks

In the last decade, many efforts have been invested into developing new methods for the local control of ion channel activity with light or chemical signals. In contrast, non-invasive mechanisms controlling ion channel expression have received substantially less attention, despite their importance in cardiac, neurological disorders and various forms of cancer. Here, we demonstrate a robust and scalable mechanism for the control of potassium channel expression on macroscopic scales in eukaryotic cell populations. As a proof-of-concept, we tested this chemically-excited mechanism in the model eukaryote to access the general suitability of our strategy for tuning ion channel activities through gene expression in other living systems. We demonstrated that a chemically-excited gene circuit robustly controls cyclic charging(hyperpolarization) of individual cells inside the microbial community by solely adjusting the expression level of the potassium channel. This “battery-mimicking” mechanism is robust to the presence of cellular noise and environmental fluctuations. Thus, this work opens new avenues in the field of synthetic biology and cellular engineering by enabling a non-invasive transformation of chemical cues into synchronized ion channel expression that modulates the electrical activity of cells on extended spatial-temporal scales. More broadly, controlling locally and globally ion channel expression holds a key to designing more effective therapies for a broad spectrum of human diseases.

## Acknowledgments

We would like to thank Dr. Luis Rubio for providing BY4741 laboratory yeast strain used in this work. This work was supported by the Programa de Atraccion de Talento 2017 (Comunidad de Madrid, 2017-T1/BIO-5654 to K.W.), Severo Ochoa (SO) Programme for Centres of Excellence in R&D from the Agencia Estatal de Investigacion of Spain (grant SEV-2016-0672 (2017-2021) to K.W. via the CBGP). In the frame of SEV-2016-0672 funding M.A. received a PhD fellowship (SEV-2016-0672-18-3: PRE2018-084946). K.W. was supported by Programa Estatal de Generacion del Conocimiento y Fortalecimiento Cientıfico y Tecnologico del Sistema de I+D+I 2019 (PGC2018-093387-A-I00) from MICIU (to K.W.). UPM Plan Propio Predoctoral fellow finances M.G.N

## Author contributions

M.G.N, S.P.G, and M.A. designed, performed most of the experiments, and analyzed data. D.R.S, contributed to plasmid and strain constructions, S.P.G and M.A, performed multi-well plate fluorescence assays. K.W designed experiments analyzed data and supervised the project. All authors contribute to the manuscript writing.

## Data and materials availability

All data are shown in the main text or the supplementary materials. Plasmids and strains are available upon request from authors.

## Materials & Methods

### 1.1 Strains and plasmid constructions

Constructs were cloned using isothermal Gibson assembly cloning. A middle-copy (∼10-30 copies) episomal plasmid pGADT7 (Takara Bio Inc.) was used to increase the concentration of proteins to buffer for the effects of intrinsic molecular noise and selected using different auxotrophic selection markers (Leucine, Uracil, and Histidine). Iacro/MarRo promoter and either standard CYC1 or ADH1 yeast terminators were cloned into activator or repressor plasmids (Fig. S2A). MarR and IacR were codon-optimized for yeast and synthesized using services delivered by Integrated DNA Technologies (IDT). The reporter plasmids include synthetic minimal promoters (synthesized with IDT) with previously identified MarR or IacR operator (31, 32) sequences upstream TATA-box and minimal CYC1 promoter and fast-degradable UBG76V-EGFP (dEGFP)(21). KcsA bacterial potassium channel was engineered to include open configuration mutations as previously described (26, 27) and modified further to include plasma membrane localization signals and c-terminus mouse ODC degron signal (33). K1 toxin was codon-optimized and synthesized throughs IDT. KcsA* and K1 toxin replace the dEGFP gene on the reporter plasmid, respectively (Fig. S2B). PCR reactions were performed using Q5 high fidelity polymerase (New England Biolabs). Correct PCR products were digested with DpnI (New England Biolabs) to remove the template and subsequently cleaned up with a DNA cleanup kit (Zymo Research) before Gibson assembly. Constructs were transformed in ultra-competent cells from E. coli DH5a strain using standard protocols. All plasmids were confirmed by colony PCR and validated with sequencing. The BY4741 laboratory yeast strain (a kind gift from Dr. Luis Rubio) carrying integrated copy constitutively expressed mCherry reporter was used to prepare competent cells and transformation of plasmids using Frozen-EZ Yeast Transformation II Kit (Zymo Research). Sequences of synthetic DNA and strains used in this study are shown in Data S1 and Table S2.

### 1.2 Multiwell plate and microscopy fluorescence measurements

Overnight culture of the yeast grown in 2% sucrose low fluorescence media (Formedium, UK) was diluted 100x and pipetted directly to a 96 well plate containing 2% sucrose and gradient of SA and IAA concentrations. Plates were incubated at 30C overnight and well mixed by shaking before performing measurements. Measurements were done with the Thermo Scientific^TM^ VarioskanTM LUX multimode microplate reader after 24h or were recorded every 10min to generate a time-lapse profile of the dEGFP and OD600. OD600 was set at an absorbance of 600nm wavelength, the fluorescence excitation and emission light at 488nm and 517nm wavelength for dEGFP. The PMP reporter activity was analyzed by measuring Thioflavin (ThT) expression using potassium clamp experiments. Overnight cultures were diluted to a total OD600 of 0.1. KCL at different concentrations was added to the diluted cultures. Aliquotes of 200 µL were pipetted from these diluted cultures to a multi-well plate containing different KCL concentrations (0 mM, 50mM, 100mM, 200mM, and 400mM) and 10uM Thioflavin T. Plates were incubated at 30 °C overnight. The next day each well was imaged in two different channels: differential interference contrast (DIC) and GFP (λEx = 488 nm; λEm = 515 nm). The image acquisition was controlled by the software µManager and Leica DMI9 and images were captured using a 10x dry objective (NA=0.32).

### 1.3 Time-lapse imaging, growth conditions, and data analysis

Live-cell imaging was performed on the Automated inverted Leica DMi8 fluorescence microscope equipped with Hamamatsu Orca Flash V3 camera that was controlled by Micro-Manager v.2.0 (https://micro-manager.org/). Images were captured with 40x dry objective NA=0.8 (Leica Inc.). Traps containing cells were imaged every 10 minutes on three different channels (DIC, GFP (Excitation: 488 Emission: 515, and mCherry Excitation: 583, Emission: 610) with CoolLed pE600 LED excitation source and standard Chroma epifluorescent filter set. Experiments were run for up to 72 hours under the continuous supply of nutrients in the microfluidic device. Acquired images were initially processed in Fiji 2.0 (https://imagej.net/Fiji) using custom scripting to extract positions with exponentially growing yeast cells. Constitutively expressed mCherry marker was used to identify exponentially growing cells and used to derive normalized dEGFP fluorescence: Dead or non-growing individuals were discarded by correcting dEGFP signal according to the formula *dEGFP*/(*dEGFP* + *mCherry*) or alternatively K1 toxin producing strain. Each image was divided into 25 regions of interest (ROIs) and analyzed separately to isolate regions where cells were actively growing and could be tracked over time. The posterior analysis was done with custom R-studio scripts. Firstly, raw data were detrended using the detrend function from “pracma” R-studio v4.0.3 package and then smoothed with Savitzky-Golay Smoothing function (savgol), from the same package, with a filter length of 15 was applied and the signal was normalized between 0 and 1 to generate heatmaps across cell traps. Amplitudes were calculated with find peaks within the Process Data using the “findpeaks” function from “pracma” R package with nups and ndowns of 6, and periods were calculated by calculating distances between successive dEGFP peaks. Phase drift was calculated by comparing time differences of successive dEGFP peaks between cell communities in a microfluidic device to derive inter-community measures. Cumulative autocorrelations traces and power spectrum densities were calculated on mean dEGFP trajectories per colony calculated for n-communities (n>20, ∼20,000 cells) using standard calculations with Matlab 2018b derived packages autocorrect and Fast Fourier Transformation (FFT). The same procedure was used to evaluate ThT fluorescence dynamics, autocorrelation function and frequency responses.

### 1.4 Mathematical model descriptions

To derive a mathematical model of the excitable circuit (Fig. 1B) we used a system of coupled ordinary differential (ODE) equations adapted from previous theoretical studies (18). Briefly, IacR and MarR protein concentrations change over time according to the following formulas:

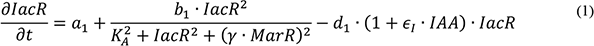

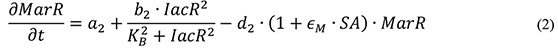

where *a1* and *a2* are basal IacR and MarR production rates. *b1* and *b2* are IacR-dependent protein production rates. *KA* and *KB* are half-max hill function coefficients, and *γ* is the rate of MarR-dependent repression. *d1* and *d2* are protein turnover rates, respectively. ϵ_*I*_ and ϵ_*M*_ are the rates of phytohormone effect on a total protein turnover. IAA and SA are modeled using square wave and sine signal generators (Matlab Inc.).

The ratio of protein turnover 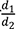 (Fig. S1A-C) represents a key bifurcation parameter that enables the transition of the system into oscillatory and excitable regimes.

To calculate nullclines we set the left-hand side of Eqs. 1 and 2 to 0 and setting ϵ_*I*_ and ϵ_*M*_ to 0.

One can derive the following terms for calculation of phase portraits MarR*(IacR) and MarR**(IacR) (Fig. S1A-C):

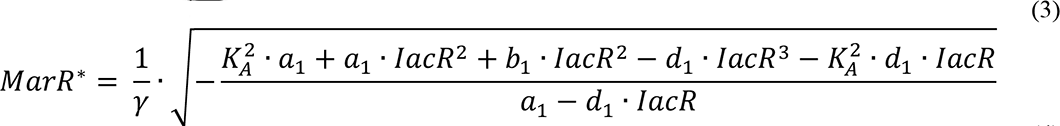

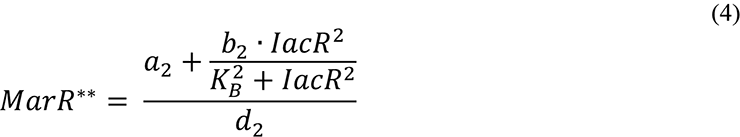

The spatial-temporal PDE model of PMP firings (Fig. S9) was simulated using space discretization dx of one colony unit. The IacR and MarR were modeled using indexed eqs. 1 and 2 *IacRi* and *MarRi* where I is yeast population index *i*=1…N , N=50.

The Hodgkin–Huxley (34) plasma membrane potential Pi for the *i*th population changes over time as follows:

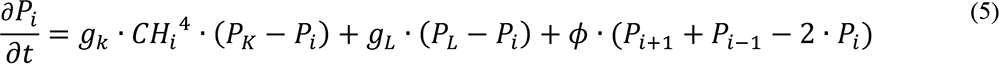

whereas the potassium channel species in the *i*th population (*CHi)*are written as:

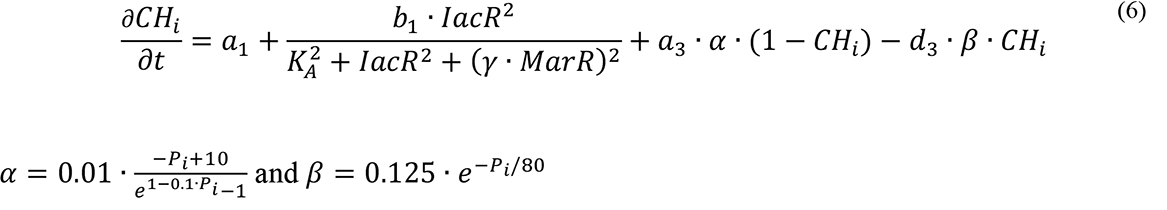

where *g*_*k*_ is electric current constant for potassium ions (channel conductance) and *g*_*L*_ is a general leak current constant, respectively. *P*_*K*_ and *P*_*L*_ are resting membrane potentials. The parameter *ϕ* denotes the rate of potassium exchange between populations that balances PMP. Parameters *a*_3_ and *d*_3_ depict activation and turnover rates of the ion channel. Model parameters are summarized in Table S1.

### 1.5 Cell loading procedure

All tubing lines were sterilized with ethanol and plugged into syringes or introduced in falcon tubes under sterile conditions. Fresh yeast colony was grown in low fluorescence media composition (Formedium, UK) with 2% sucrose as a carbon source or 2% glucose (K1 toxin-producing strain). The next day, yeast cultures were diluted 10-20 times approximately to 0.2-0.4 O.D600 to obtain highly concentrated cells that were transferred to a 50mL falcon tube for loading. 60mL media syringes were filled with 25mL of inducing media (2% sucrose + 0.5%galactose) with or without compounds and 50mL waste falcons tubes were filled with 10mL of DDI water. Before loading, devices were vacuumed for at least 20 minutes to remove all the air from the channels and traps. Syringes and falcon tubes were placed on the height control system and lines were connected as follows; media syringes were plugged first and kept above all other inputs to prevent media contamination. Adjusting the height of the cells-containing falcon tube as well as media and waste aids in controlling the cell seeding in the traps. Although many cells pass through the chip directly towards the waste port, few cells got captured via microvalves and seeded the traps. Once 10-20 cells were captured in each trapping region, the flow from the cell loading port was reverted by decreasing the height to the same level as for the auxiliary waste.

### 1.6 Microfluidic Mold fabrication

Molds for the production of microfluidic device (16) were designed in Inkscape and printed on plastic sheets with the monochrome laser printer at 1200dpi resolution as described in (16). A density of Ink deposition was used to control the feature height. Plastic wafers were cut and transferred to the thermal oven set to 160°C to shrink by one-third of the original size, then baked again for 10 min to smoothen and harden the ink. Finally, molds were cleaned with soap, rinsed with isopropanol and DDI water and dried using a nitrogen gun, and secured with Scotch tape before use.

### 1.7 Soft Lithography

Molds were introduced in plastic 90mm Petri dishes and fixed with double-sided tape. Dowsil Sylgard 184 Polydimethylsiloxane (PDMS) was properly mixed in a 10:1 (w/w) ratio of elastomer and curing agent and stirred until the uniform consistency was achieved. Approximately 27mL of the homogeneous mixture was poured into each petri dish and completely degassed using the 8

CFM 2-stage vacuum pump for approximately 20 minutes. Degassed PDMS was cured at 80oC for 2h. Cured PDMS was removed from the petri dish, separated from the wafer, and cut to extract the individual chips. Fluid access ports were punched with a 0.7mm diameter World Precision Instruments (WPI) biopsy puncher and flushed with ethanol to remove any remaining PDMS. Individual chips were cleaned with ethanol and DDI water and Scotch tape to remove any remaining dirt particles.

### 1.8 Microfluidic device bonding

At least one day before use, individual chips and coverslips were cleaned in the sonic bath and rinsed in ethanol, isopropanol, and water. Both surfaces were exposed to Corona SB plasma treater (ElectroTechnics Model BD-20AC Hand-Held Laboratory Corona Treater) between 45 seconds to 1 minute, then surfaces were brought together and introduced at 80°C in an oven overnight to obtain the enhanced bond strength.

## Supplementary Information

**Fig. S1.**
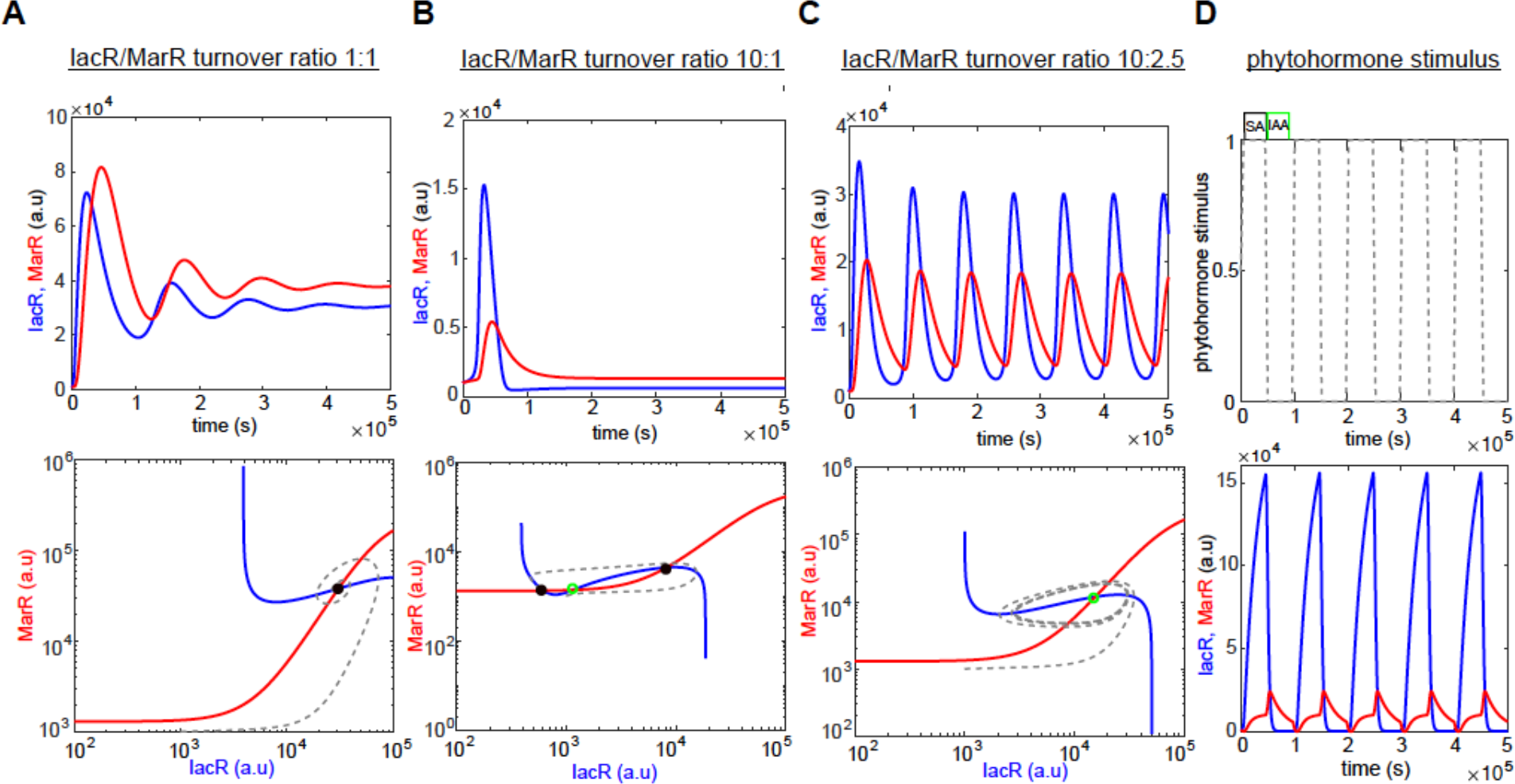
Computer model simulations of IacR-MarR feedback circuit reveal excitable dynamics controlled by phytohormones. **(A-C)** Computer simulations of a dual-feedback circuit model (upper panel) and corresponding phase portraits with nullclines (lower panel) for the different ratio of IacR-MarR receptor turnover (1:1 (A), 10:1 (B), 10:2 (C)). Stable states are marked with black-filled circles and unsteady states with a green empty circle, respectively. Note that a higher turnover ratio produces the transition into an excitable regime around the unstable state(B). **(D)** Environmental phytohormone stimuli push the system into an excitable regime by directly increasing and successively decreasing the IacR-MarR turnover ratio(B).

**Fig. S2.**
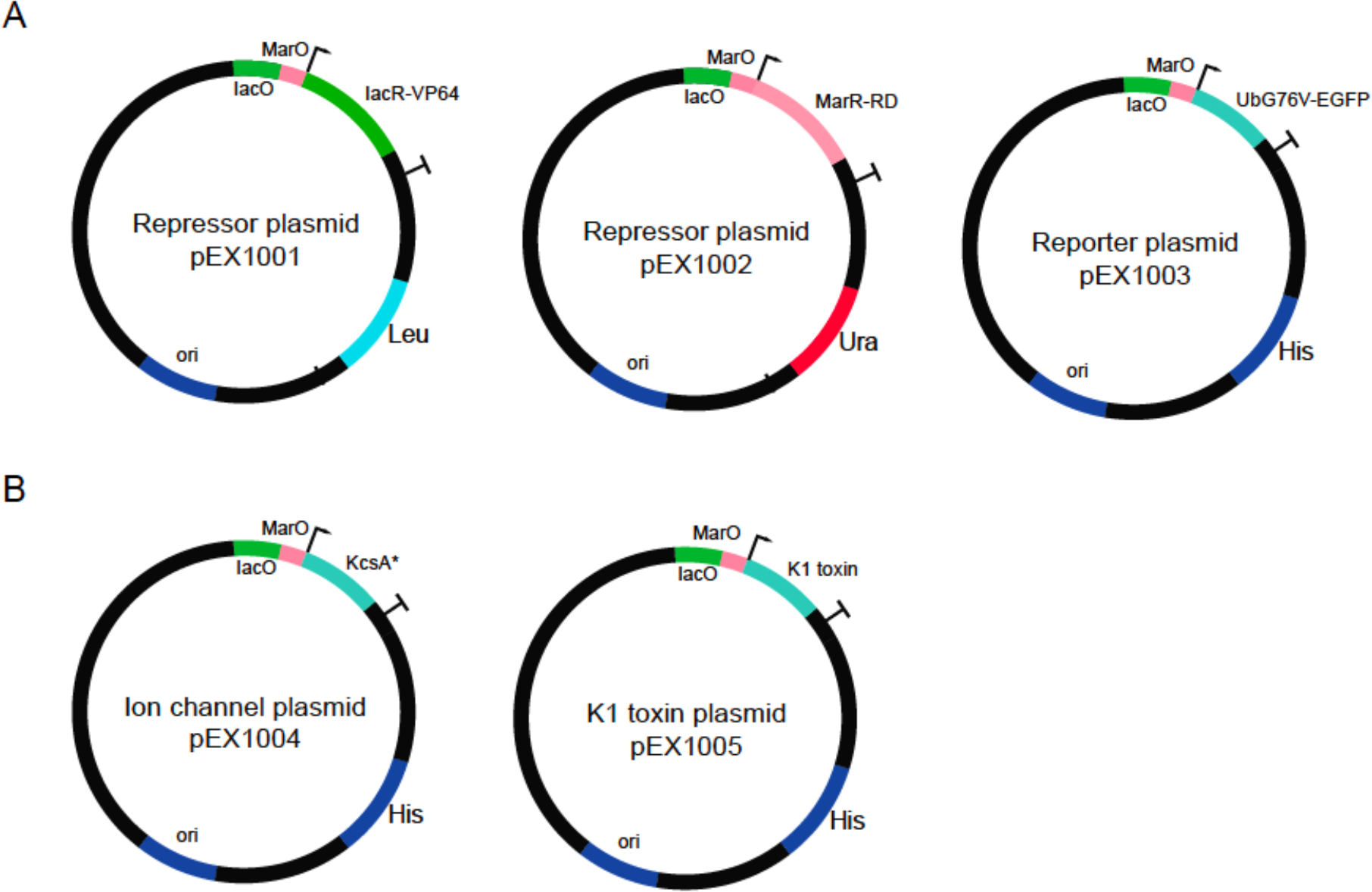
The plasmids used in the study. **(A, B)** Plasmid map schematics were used in the study. (A) Activator (IacR) and repressor (MarR) and reporter (dEGFP) plasmids were used to build an excitable circuit as shown in Fig. 1B. (B) Additional plasmids are used to construct an electrical signaling mediator; an engineered ion channel (KcsA*) or host interface (K1 toxin).

**Fig. S3.**
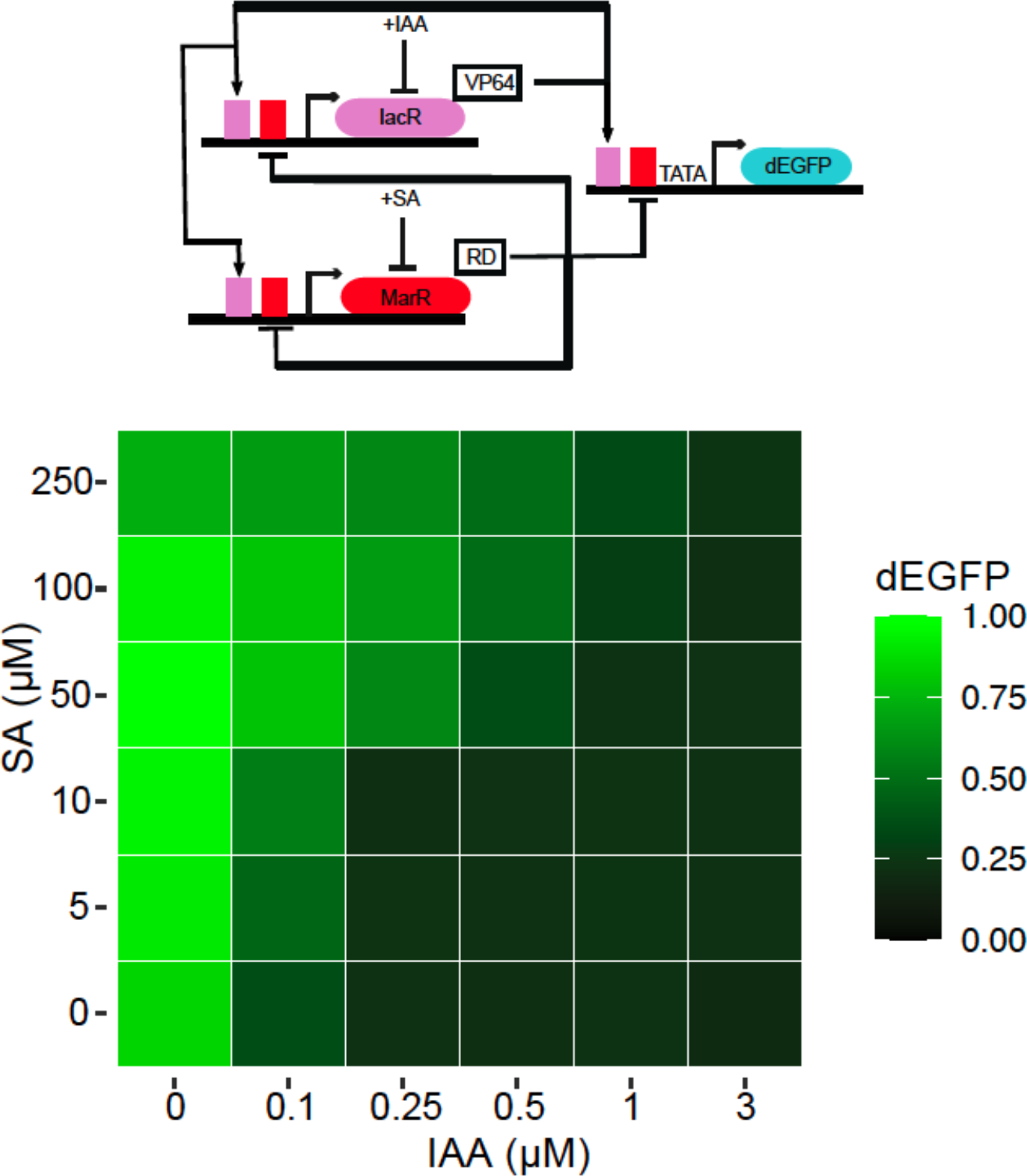
Precise control of circuit output with dual phytohormones. A heat map of excitable circuit response recorder with dEGFP reporter (upper panel) to chemical gradients (lower panel). Note a precise control of circuit output is possible by using a dual phytohormone strategy. Mean measurements from three biological replicates (n=3) are shown.

**Fig. S4.**
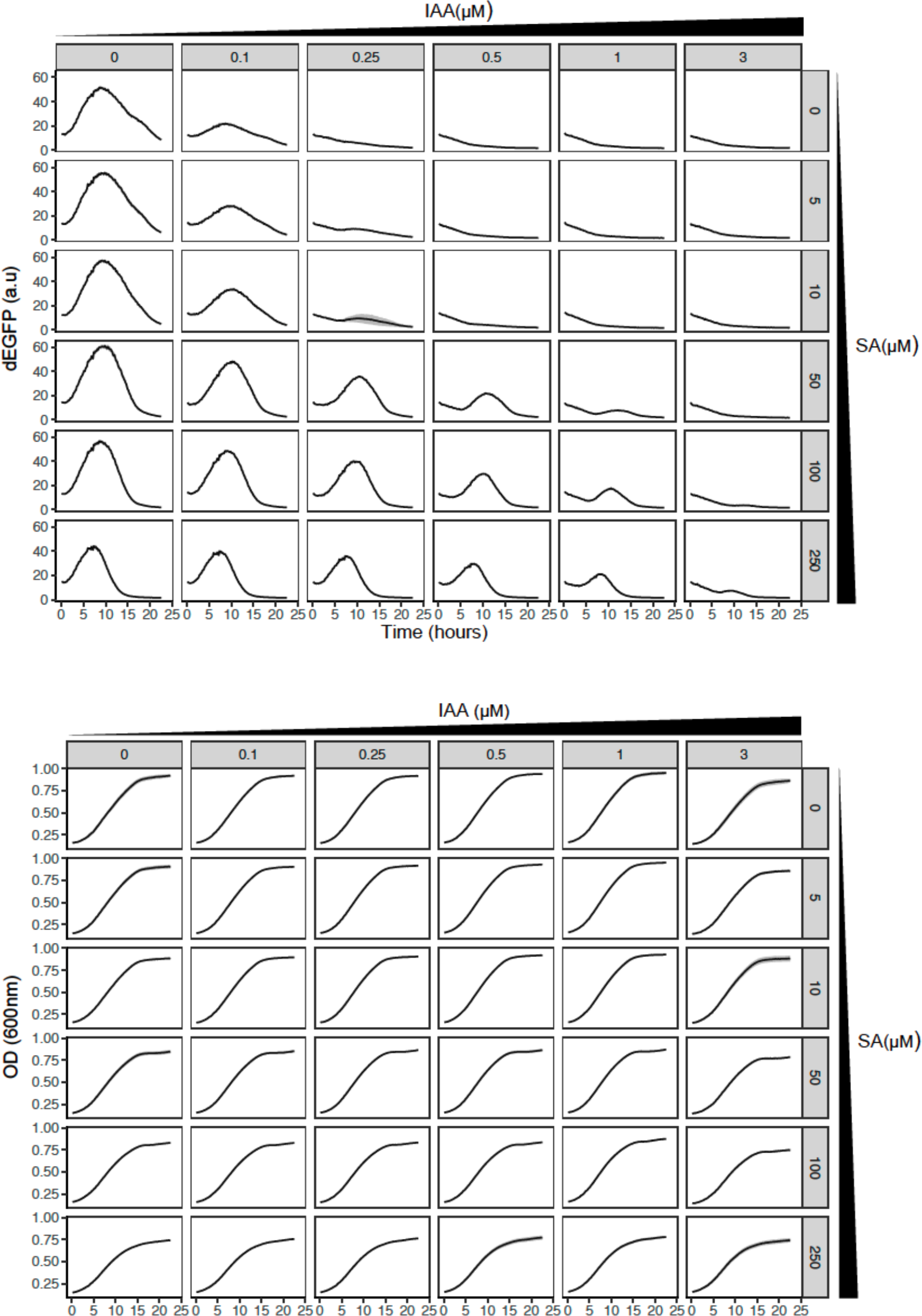
Time-lapse synthetic circuit characterization using high-throughput fluorescence assays. dEGFP fluorescence normalized to OD600 (upper panel) and OD600 absorbance (lower panel) for 24-hour measurements using graded changes of phytohormones was used to analyze the excitable potential of the synthetic circuit. Mean measurements with variations(shaded regions) from three biological replicates (n=3) are shown.

**Fig. S5.**
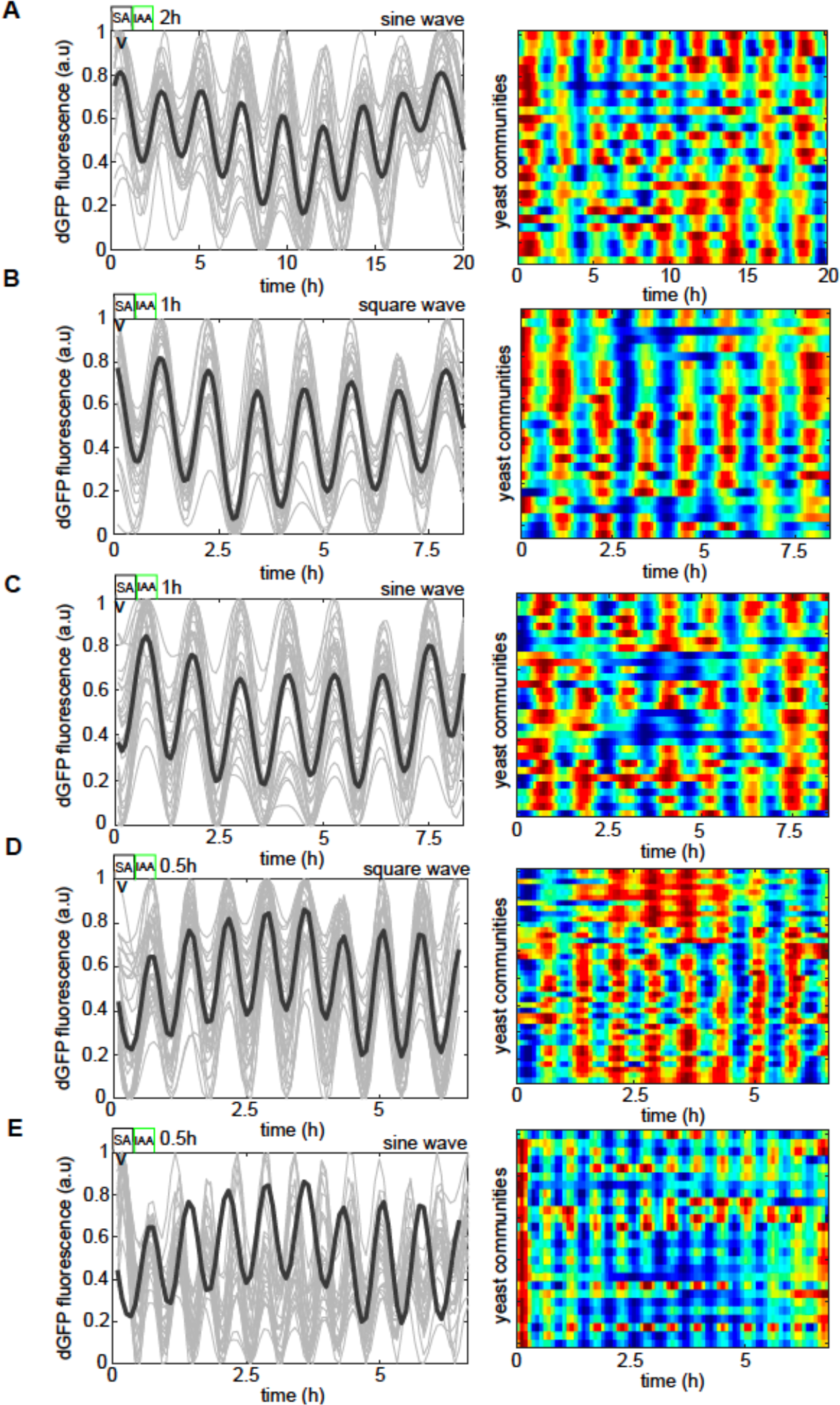
The frequency of phytohormone stimuli determines the excitable dynamics of the reporter gene under different types of environmental drivers. **(A-E)** Left panel, dEGFP time evolutions from all traps recorded in the microfluidic device. Right panel, the heat map of dEGFP fluorescence for each of the recorded trapping regions corresponds to time traces(left panel). (A), Periodic 2-hour phytohormone wave of sine shape, (B, C) 1-hour periodic waves of square(B) and sine (C) shapes. (D, E) 30 min periodic wave of square (D) and sine (E) shapes, respectively. Note that the frequency of dEGFP firings mirrors exactly that of the chemical microenvironment. Color coding is as in Fig. 1.

**Fig. S6.**
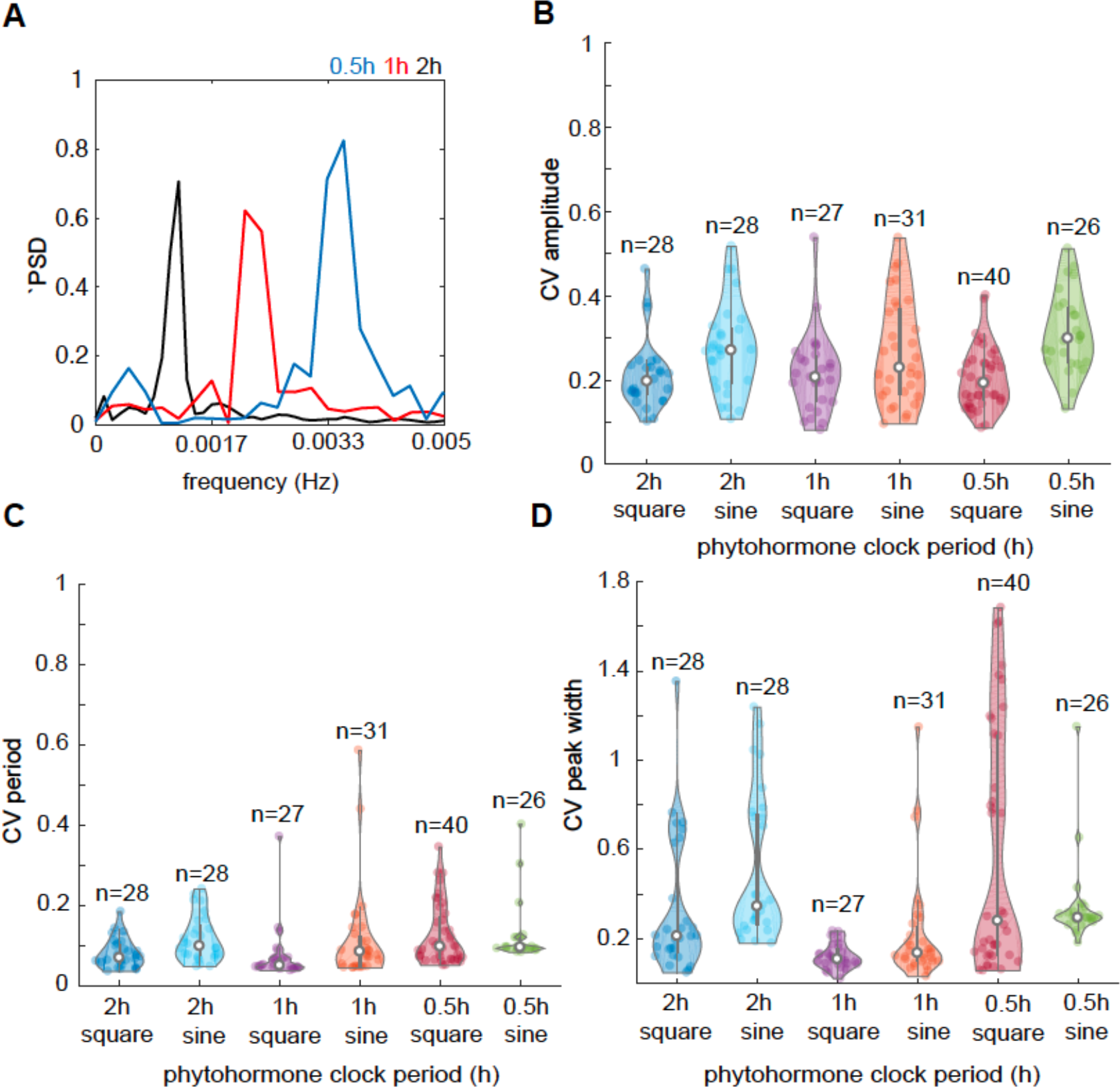
The excitable circuit shows low variation in the period, amplitude, and response peak width regardless of the stimulation period. **(A)** dEGFP Frequency spectra (PSD) of globally coordinated populations for different periods of phytohormone rhythms (2h, 1h, and 30min respectively). (**B-D**) Coefficients of variation (CV) in the amplitude(B), periods(C), and response peak width(D) of dEGFP fluorescent reporter under different environments and stimulus periods. Note generally very low CV (CV < 1) indicates low variation in dEGFP reporter signal characteristics.

**Fig. S7.**
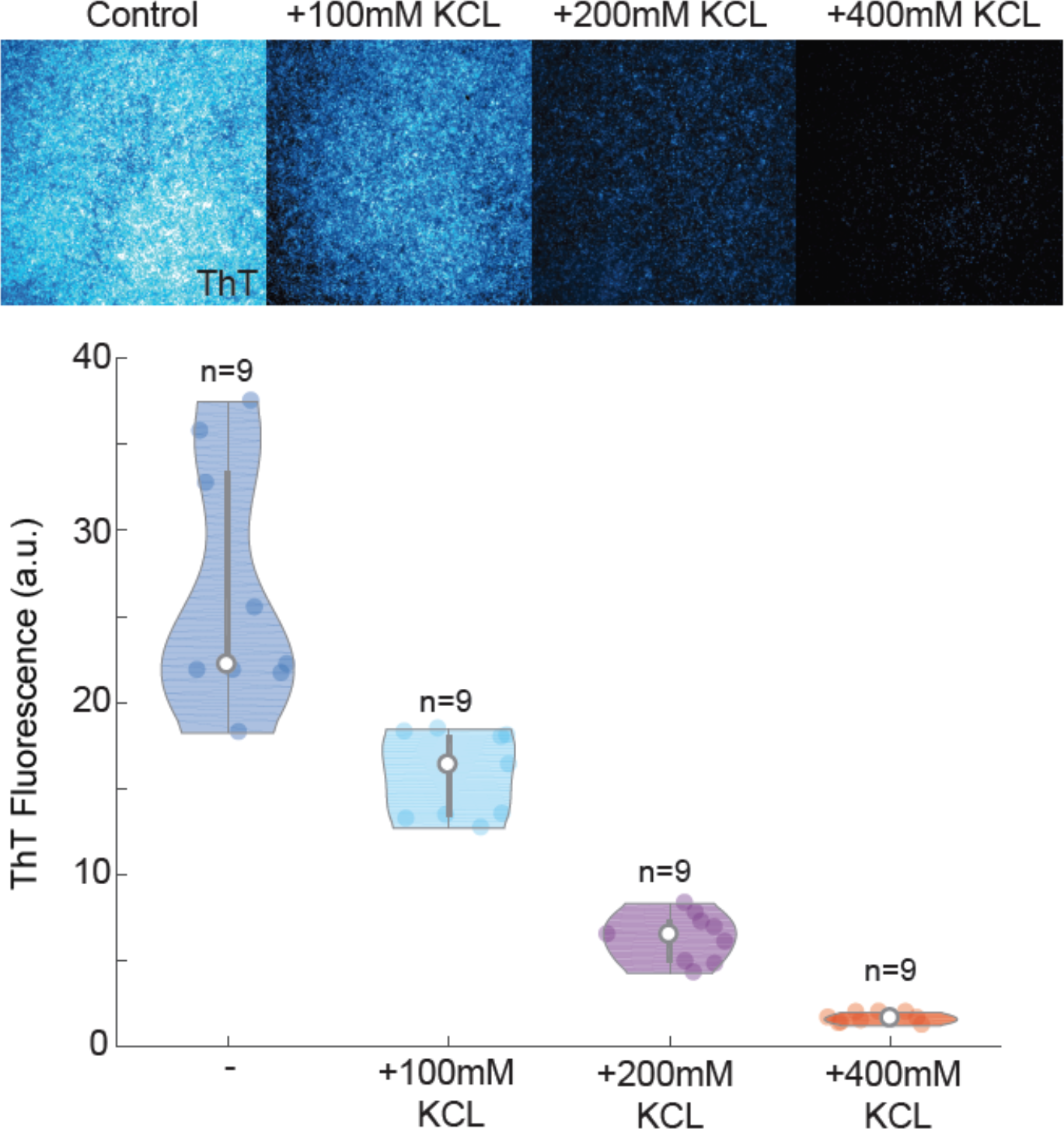
Testing ThT reporter activity in yeast cells through potassium clamp experiments. Upper panel, screenshots from microscopy imaging of yeast after 24h in different KCL concentrations, and quantification of mean ThT fluorescence (lower panel) in independent biological replicates (n=9). Note, a near-linear drop in THT fluorescence after excessive amounts of potassium have been applied.

**Fig. S8.**
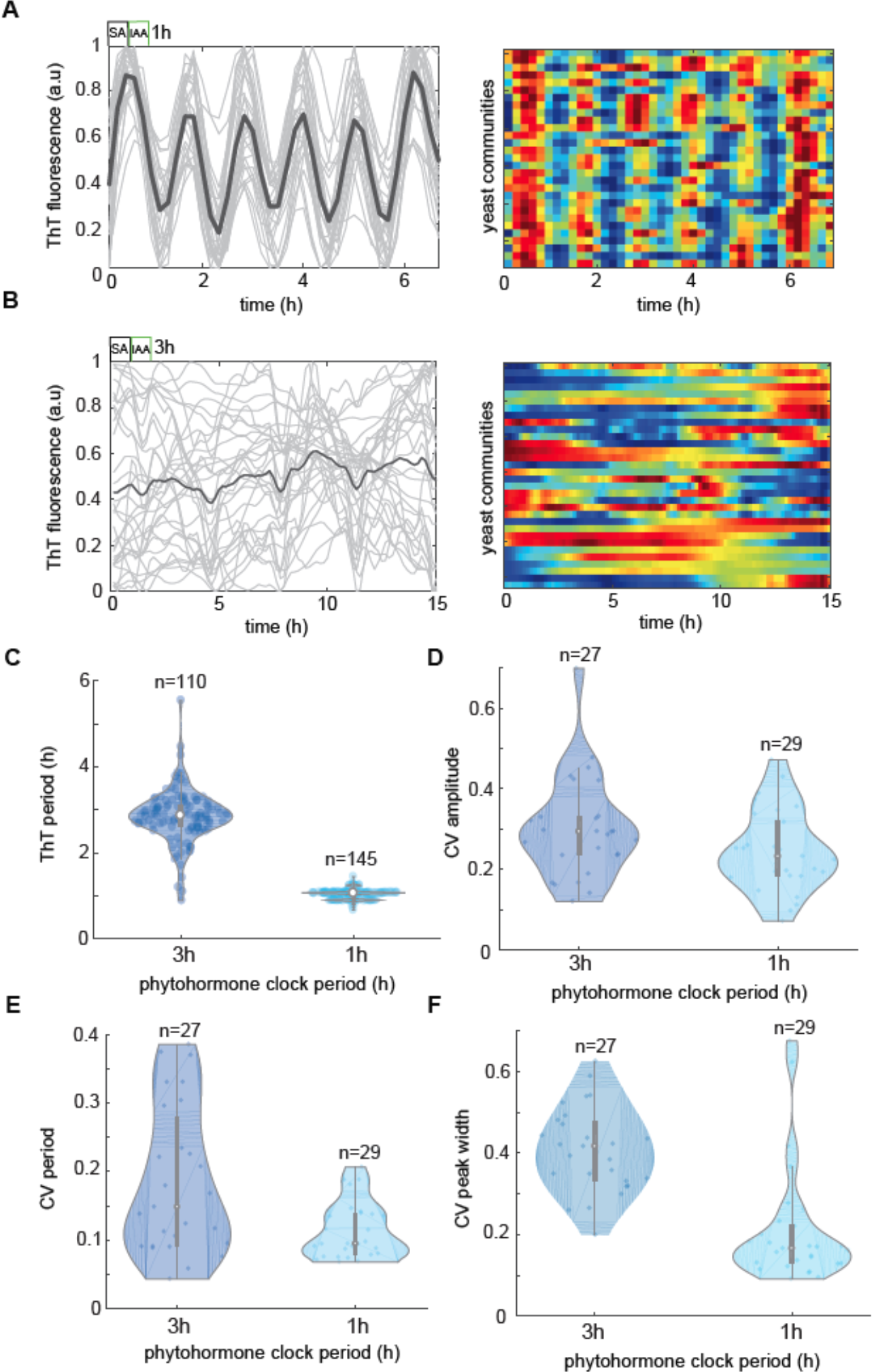
PMP firings are consistent with the environmental period of phytohormone stimuli only in the cells with the excitable circuit. **(A)** Left panel, ThT fluorescence time evolutions from all traps recorded in the microfluidic device. Right panel, the heat map of ThT fluorescence for each of the recorded trapping regions corresponds to time traces (left panel) for 1-hour stimulation with antithetic IAA and SA pulses. (**B**) ThT dynamics under the 3-hour environmental stimulus (left, time course and right, heat map) in the control strain populations that lack the excitable circuit. Color coding as in Fig. 1**. (C)** Observed the ThT signal period in the function of the phytohormone stimulation period. **(D-F)** Coefficients of variation (CV) in the amplitude(D), periods(E), and response peak width(D) of dEGFP fluorescent reporter under different environments and stimulus periods. Note generally very low CV (CV < 1) indicates low variation in general response characteristics.

**Fig. S9.**
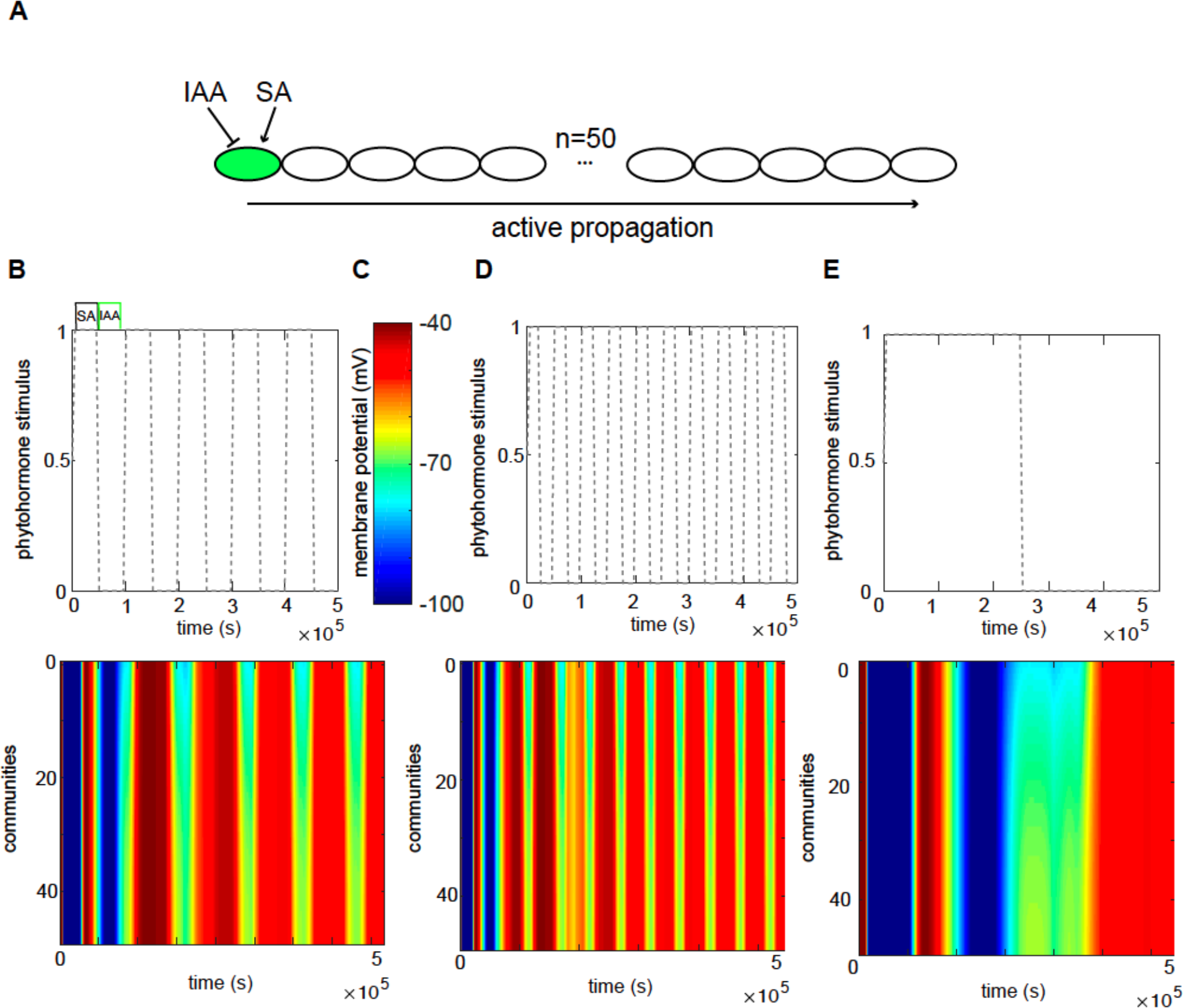
A spatial-temporal computer model predicts the active electrical signal propagation dictated by dynamic changes in ion channel abundance with the environmental pulse. **(A)** Schematics of model simulation setup. The model included the 50 adjacent populations of yeast colonies. The leftmost colony is stimulated with high amplitude pulses of IAA and SA that could trigger active signal propagation by PMP changes. (**B-E**) Phytohormone stimulus function (upper panel) and predicted PMP changes (heat map, lower panel) across populations for intermediate (B), fast (D), and slow (E) phytohormone changes in the environment. (C) color coding for heat maps.

**Table S1.**
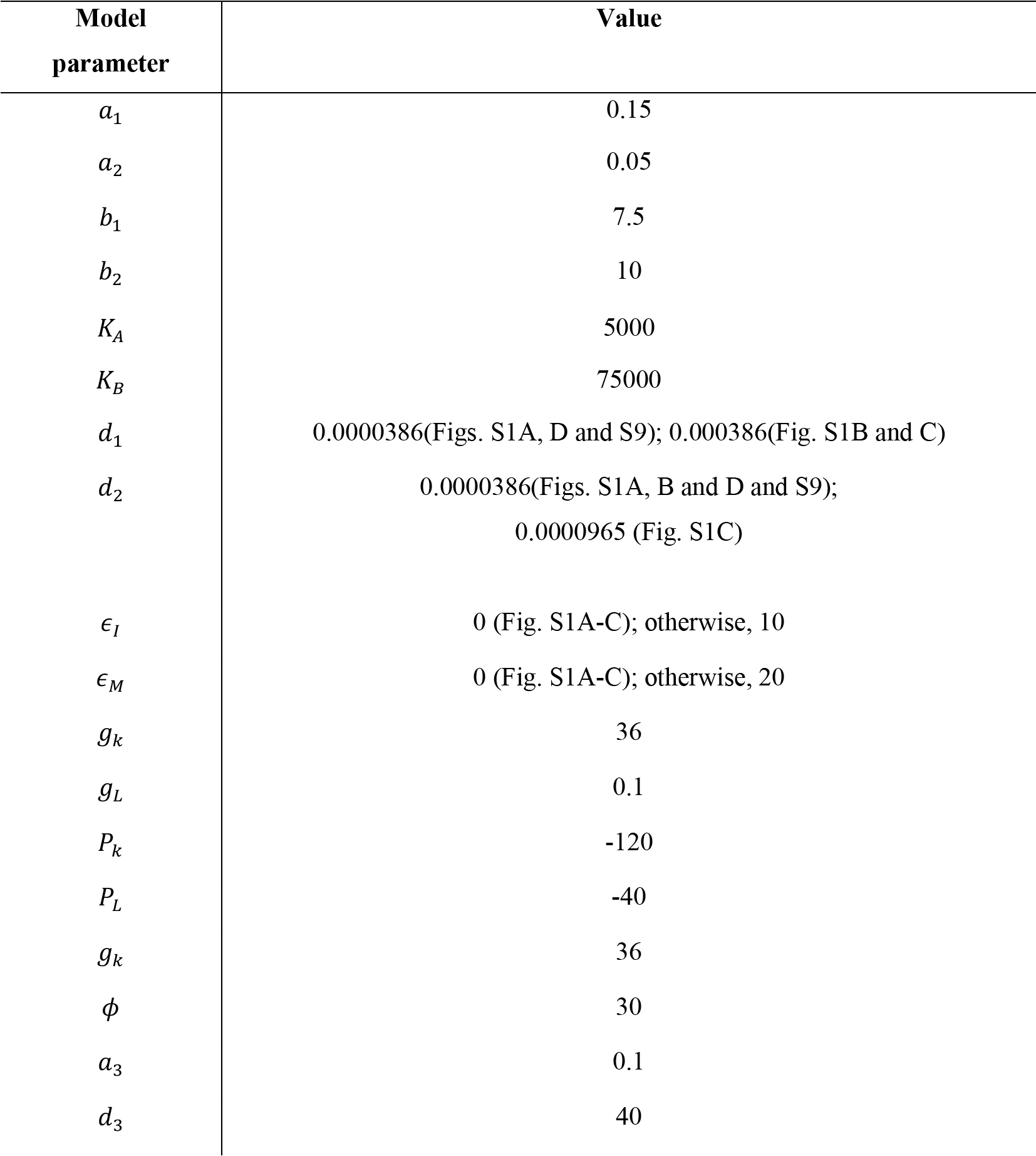
Model parameters.

**Table S2.** Yeast strains.

| Strain name | Parental strain | Composition | Markers/autotrophy |
| --- | --- | --- | --- |
| sRedM | BY4741 MATa<br>his3Δ1 leu2Δ0<br>met15Δ0 ura3Δ0 | BY4741 pGK1:: mCherry | KanMX (G418) |
| sEXGFP | sRedM | sRedM<br>pMarIac:: IacR-VP64 (pEX1001)<br>pMarIac::MarR-VP64 (pEX1002)<br>pMarIac::dEGFP(pEX1003) | KanMX (G418)<br>Leu, Ura , His |
| sPMPEX | BY4741 MATa<br>his3Δ1 leu2Δ0<br>met15Δ0 ura3Δ0 | BY4741<br>pMarIac:: IacR-VP64 (pEX1001)<br>pMarIac::MarR-VP64 (pEX1002)<br>pMarIac::KcsA* (pEX1004) | Leu, Ura , His |
| sEmpty(control/<br>sensitive strain) | BY4741 MATa<br>his3Δ1 leu2Δ0<br>met15Δ0 ura3Δ0 | Empty plasmids | Leu, Ura , His |
| sKiller | sRedM | sRedM<br>pMarIac:: IacR-VP64 (pEX1001)<br>pMarIac::MarR-VP64 (pEX1002)<br>pMarIac::K1 | KanMX (G418)<br>Leu, Ura , His |
|  |  | (pEX1005) |  |

Movie S1.

Excitable gene circuit shows coordinated oscillations. Related to Fig 1. Left Panel, Differential interference contrast (DIC), middle panel, dEGFP(cyan) and mCherry(cell marker), and right panel, dEGFP color coded in heat map colors (blue(low signal), red(high signal)) of the sampled yeast colony grow in microfluidics device with 2-hour cycles of phytohormone stimuli.

Movie S2.

PMP oscillations controlled by phytohormone rhythms (3h stimuli). Related to Fig 2. Left Panel, Differential interference contrast (DIC), middle panel, THT(cyan) and), and right panel, ThT color coded in heat map colors (blue(low signal), red(high signal)) of the sampled yeast colony grow in microfluidics device with 3-hour cycles of phytohormone stimuli.

Movie S3.

PMP oscillations controlled by phytohormone rhythms (1h stimuli). Related to Fig 2. Left Panel, Differential interference contrast (DIC), middle panel, THT(cyan) and), and right panel, ThT color coded in heat map colors (blue(low signal), red(high signal)) of the sampled yeast colony grow in microfluidics device with 1-hour cycles of phytohormone stimuli.

Movie S4.

The absence of PMP oscillations in the control strain. Related to Fig. S8B. Left Panel, Differential interference contrast (DIC), middle panel, THT(cyan) and), and right panel, ThT color coded in heat map colors (blue(low signal), red(high signal)) of the sampled yeast colony grow in microfluidics device with 3-hour cycles of phytohormone stimuli.

Movie S5.

PMP oscillations in sensitive strain controlled by phytohormone rhythms in cocultures. Related to Fig. 3. Left Panel, Differential interference contrast (DIC), right panel, THT in sensitive strain (cyan) and (killer strain, mCherry of the sampled yeast two strain cocultures grow in microfluidics device with 3-hour cycles of phytohormone stimuli.

**Dataset S1.**
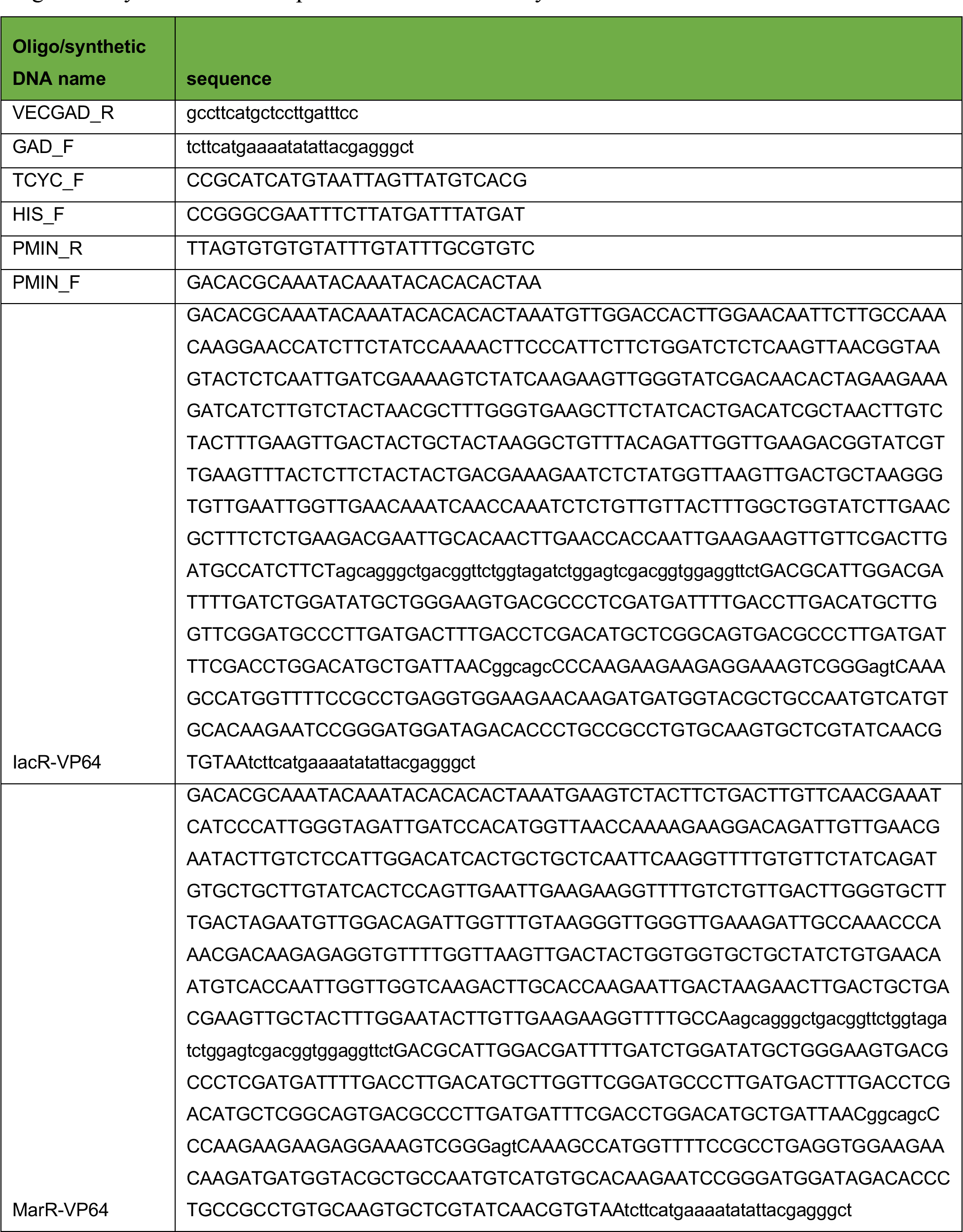
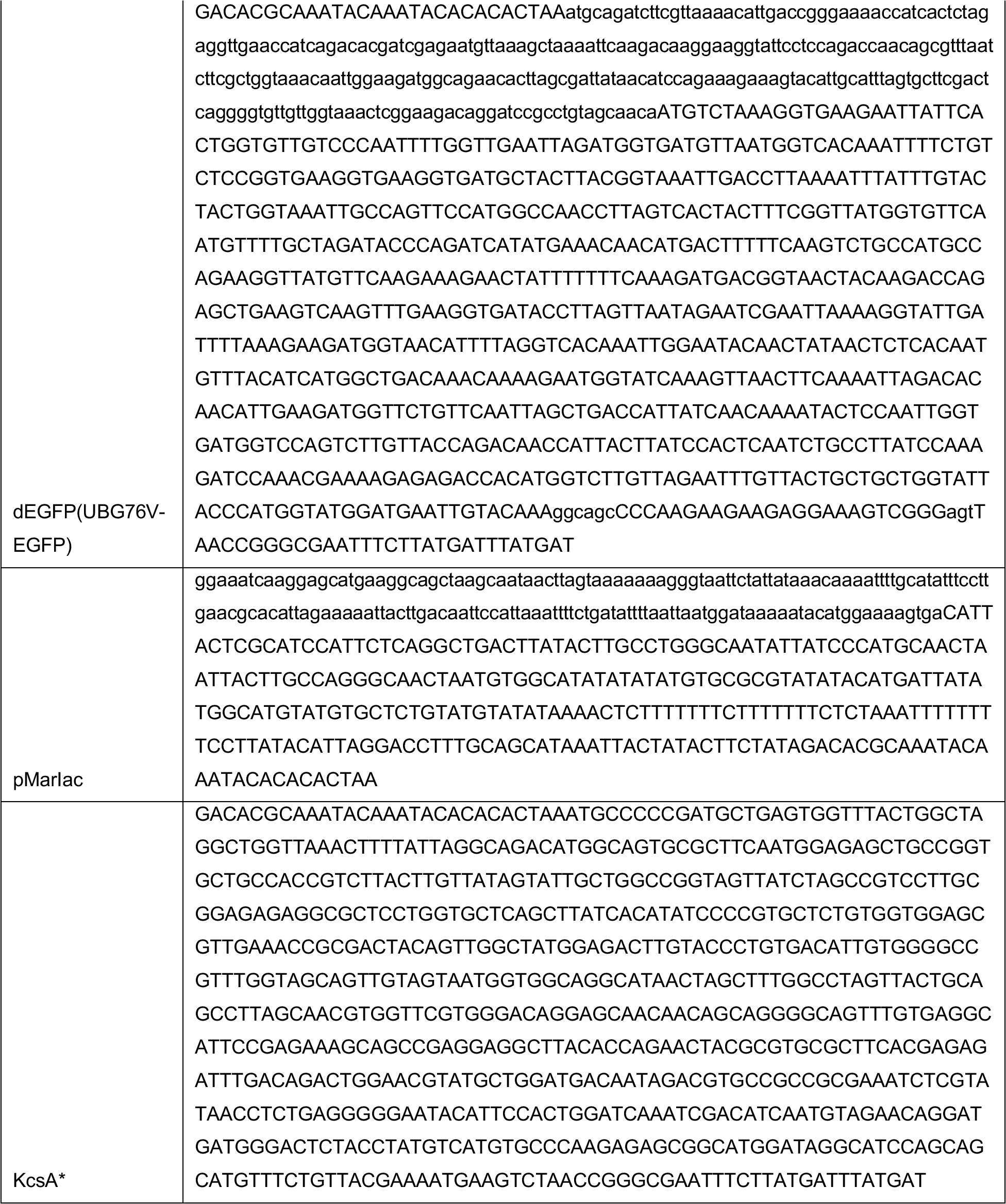
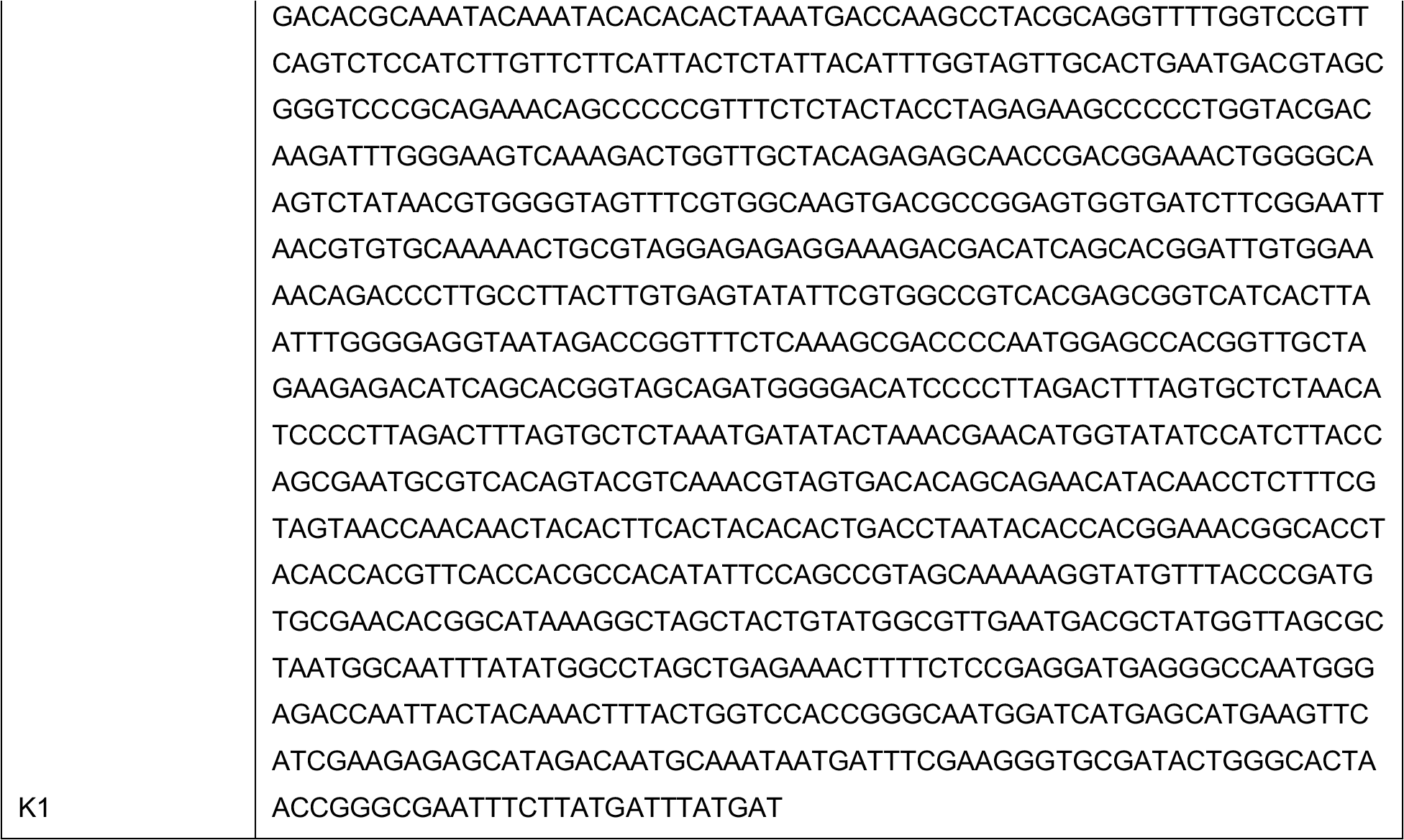
Oligos and synthetic DNA sequences used in this study.

